# Analysis of *Brevundimonas subvibrioides* developmental signaling systems reveals unexpected differences between phenotypes and c-di-GMP levels

**DOI:** 10.1101/531418

**Authors:** Lauryn Sperling, Volkhard Kaever, Patrick D. Curtis

## Abstract

The DivJ-DivK-PleC signaling system of *Caulobacter crescentus* is a signaling network that regulates polar development and the cell cycle. This system is conserved in related bacteria, including the sister genus *Brevundimonas*. Previous studies had shown unexpected phenotypic differences between the *C. crescentus divK* mutant and the analogous mutant of *Brevundimonas subvibrioides*, but further characterization was not performed. Here, phenotypic assays analyzing motility, adhesion, and pilus production (the latter characterized by a newly discovered bacteriophage) revealed that *divJ* and *pleC* mutants have mostly similar phenotypes as their *C. crescentus* homologs, but *divK* mutants maintain largely opposite phenotypes than expected. Suppressor mutations of the *B. subvibrioides divK* motility defect were involved in cyclic-di-GMP (c-di-GMP) signaling, including the diguanylate cyclase *dgcB*, and *cleD* which is hypothesized to affect flagellar function in a c-di-GMP dependent fashion. However, the screen did not identify the diguanylate cyclase *pleD.* Disruption of *pleD* in *B. subvibrioides* caused hypermotility in wild-type, but not in the *divK* background. Analysis of c-di-GMP levels in these strains revealed incongruities between c-di-GMP levels and displayed phenotypes with a notable result that suppressor mutations altered phenotypes but had little impact on c-di-GMP levels in the *divK* background. Conversely, when c-di-GMP levels were artificially manipulated, alterations of c-di-GMP levels in the *divK* strain had minimal impact on phenotypes. These results suggest that DivK performs a critical function in the integration of c-di-GMP signaling into the *B. subvibrioides* cell cycle.

**Importance:** Cyclic-di-GMP signaling is one of the most broadly conserved signaling systems in bacteria, but there is little understanding of how this system directly affects the physiology of the organism. In *C. crescentus*, c-di-GMP has been integrated into the developmental cell cycle, but there is increasing evidence that environmental factors can impact this system as well. The research presented here suggests that developmental signaling could impact physiological Processes in c-di-GMP dependent and independent ways. This hints that the integration of these signaling networks could be more complex than previously hypothesized, which could have a bearing on the larger field of c-di-GMP signaling. In addition, this work further examines how much models developed in one organism can be extrapolated to related organisms.

## Introduction

Though model organisms represent a small portion of the biodiversity found on Earth, the research that has resulted from their study shapes much of what we know about biology today. The more closely related species are to a model organism, the more that theoretically can be inferred about them using the information from the model organism. Modern genomic studies have given this research an enlightening new perspective. Researchers can now compare the conservation of particular systems genetically. Using model organisms can be a very efficient and useful means of research, but the question still remains of how much of the information gained from the study of a model can be extrapolated unto other organisms. Though genomic comparison shows high levels of conservation between genes of different organisms, this does not necessarily mean the function of those genes or systems has been conserved. This phenomenon seems to be evident in the *Caulobacter crescentus* system.

*C. crescentus* is a Gram-negative alphaproteobacterium that lives a dimorphic lifestyle. It has been used as a model organism for the study of cell cycle regulation, intracellular signaling, and polar localization of proteins and structures in bacteria. The *C. crescentus* life cycle begins with the presynthetic (G_1_) phase in which the cell is a motile “swarmer cell” which contains a single flagellum and multiple pili at one of the cell’s poles [for review, see (1)]. During this period of the life cycle, the cell cannot replicate its chromosome or perform cell division. Upon differentiation, the cell dismantles its pili and ejects its flagellum. It also begins to produce holdfast, an adhesive polysaccharide, at the same pole from which the flagellum was ejected. The cell then develops a stalk, projecting the holdfast away from the cell at the tip of the stalk. The differentiation of the swarmer cell to the “stalked cell” marks the beginning of the synthesis (S) phase of the cell life cycle as chromosome replication is initiated. As the stalked cell replicates its chromosome and increases its biomass in preparation for cell division, it is referred to as a predivisional cell. Toward the late predivisional stage, it again becomes replication incompetent and enters the postsynthetic (G_2_) phase of development. At the end of the G_2_ phase, the cell completes division forming two different cell types. The stalked cell can immediately reenter the S phase, while the swarmer cell moves once again through the G_1_ phase.

*Brevundimonas subvibrioides* is another Gram-negative alphaproteobacterium found in oligotrophic environments that lives a dimorphic lifestyle like that of *C. crescentus*. *Brevundimonas* is the next closest genus phylogenetically to *Caulobacter*. According to a Pairwise Average Nucleotide Identity (ANI) test, their genomes are approximately 74% identical. Bioinformatic analyses showed that all developmental signaling proteins found in the *C. crescentus* cell cycle are conserved *B. subvibrioides* (2, 3). However, little physiological characterization has been performed. Conservation of genes does not necessarily mean conservation of function or properties (3). Essential gene studies within the Alphaproteobacteria have shown that gene essentiality/non-essentiality in one organism does not always correspond with that in another organism (3–6). Analyses that have been performed on *C. crescentus* and *B. subvibrioides* have shown many similarities in gene essentiality between the two, but have shown several surprising differences as well (3).

In *C. crescentus*, the DivJ-DivK-PleC system controls the spatial activation of one of the master regulators in *C. crescentus*, CtrA (1, 7). This system is a prime example of how *C. crescentus* has evolved traditional two-component proteins into a more complex signaling pathway and, as a result, has developed a more complex life cycle. The DivJ-DivK-PleC pathway consists of two histidine kinases (PleC and DivJ) and a single response regulator (DivK) (8, 9). DivJ is absent in swarmer cells but is produced during swarmer cell differentiation. It then localizes to the stalked pole (8). DivJ is required for, among other things, proper stalk placement and regulation of stalk length (8). *C. crescentus divJ* mutants display filamentous shape, a lack of motility, and holdfast overproduction (8, 9).

PleC localizes to the flagellar pole during the predivisional cell stage (10). Though structurally a histidine kinase PleC acts as a phosphatase, constitutively de-phosphorylating DivK (8, 9). *C. crescentus pleC* mutants display a lack of pili, holdfast, and stalks, and have paralyzed flagella leading to a loss of motility (11–13). DivK is a single-domain response regulator (it lacks an output domain) whose location is dynamic throughout the cell cycle (9, 14). DivK remains predominantly unphosphorylated in the swarmer cell, while it is found mostly in its phosphorylated form in stalked cells. Photobleaching and FRET analysis show that DivK shuttles rapidly back and forth from pole to pole in the pre-divisional cell depending on its phosphorylation state (9). Previous studies have shown that phosphorylated DivK localizes bipolarly while primarily unphosphorylated DivK is delocalized throughout the cell (9). A *divK* cold-sensitive mutant suppresses the non-motile phenotype of *pleC* at 37°C. However, at 25°C, it displays extensive filamentation much like the *divJ* mutant (15). Additionally, filamentous *divK* mutants sometimes had multiple stalks, though the second stalk was not necessarily polar (15). Furthermore, electron microscopy of *divK* disruption mutants led to the discovery that they lack flagella (15).

Upon completion of cytokinesis, PleC and DivJ are segregated into different compartments, thus DivK phosphorylation levels in each compartment are dramatically different. This leads to differential activation of CtrA in the different compartments (9, 16). In the swarmer cell, the de-phosphorylated DivK leads to the downstream activation of CtrA. CtrA in its active form binds the chromosome at the origin of replication and prevents DNA replication (17, 18). The opposite effect is seen in stalked cells where highly phosphorylated DivK results in the inactivation of CtrA and, therefore, permits DNA replication (19).

Gene essentiality studies in *B. subvibrioides* led to the discovery of a discrepancy in the essentiality of DivK. In *C. crescentus* DivK is essential for growth, while in *B. subvibrioides* DivK is dispensable for growth (3, 15). Further characterization found dramatic differences in the phenotypic consequences of disruption. Through the use of a cold-sensitive DivK allele or by ectopic depletion, *C. crescentus divK* disruption largely phenocopies *divJ* disruption in cell size and motility effects (8, 9, 15). This is to be expected as DivK∼P is the active form and both *divJ* or *divK* disruption reduce DivK∼P levels. In *B. subvibrioides*, disruption of *divJ* leads to the same effects in cell size, motility, and adhesion (3). However, *divK* disruption leads to opposite phenotypes of cell size and adhesion, and while motility is impacted it is likely by a different mechanism.

While the previous study revealed important differences between the organisms, it did not analyze the impact of PleC disruption, nor did it examine pilus production or subcellular protein localization. The work presented here further characterizes the DivJ-DivK-PleC signaling system in *B. subvibrioides* and begins to address the mechanistic reasons for the unusual phenotypes displayed by the *B. subvibrioides divK* mutant.

## Materials and Methods

### Strains and growth conditions

A complete list of strains used in this study is presented in the appendix (see Table 1). *Brevundimonas* strains were cultured at 30°C on PYE medium (2 g peptone, 1 g yeast extract, 0.3 g MgSO_4_ · 7H_2_O, 0.735 CaCl_2_) (20). Kanamycin was used at 20 µg/ml, gentamycin at 5 µg/ml, and tetracycline at 2 µg/ml when necessary. PYE plates containing 3% sucrose were used for counter-selection. *Escherichia coli* was cultured on Luria-Bertani (LB) medium (10 g/L tryptone, 10 g/L NaCl, 5 g/L yeast extract) at 37°C. Kanamycin was used at 50 µg/ml, gentamycin at 20 µg/ml, and tetracycline at 12 µg/ml when necessary.

### Mutant generation

The *B. subvibrioides ΔdivJ, ΔdivK,* and *ΔdivJΔdivK* mutants were used from a previous study (3). The *B. subvibrioides ΔpleC* construct was made by PCR amplifying an upstream fragment of ∼650 bps using primers PleC138Fwd (attgaagccggctggcgccaCCAGATCGAAAAGGTGCAGCCC) and PleCdwRev (tctaggccgcGCCCCGCAAGGCGCTCTC) and a downstream fragment of ∼550 bps using primers PleCupFwd (cttgcggggcGCGGCCTAGAGCCGGTCA) and PleC138Rev (cgtcacggccgaagctagcgGGTGCTGGGATGAAGACACG). The primers were designed using the NEBuilder for Gibson Assembly tool online (New England Biolabs) and were constructed to be used with the pNPTS138 vector (MRK Alley, unpublished). Following a digestion of the vector using HindIII and EcoRI the vector along with both fragments were added to Gibson Assembly Master Mix (New England Biolabs) and allowed to incubate for an hour at 50°C. Reactions were then transformed into *E. coli* and correct plasmid construction verified by sequencing to create plasmid pLAS1. This plasmid was used to delete *pleC* in *B. subvibrioides* as previously described (3).

To create insertional mutations in genes, internal fragments from each gene were PCR amplified. A fragment from gene *cpaF* was amplified using primers cpaFF (GCGAACAGAGCGACTACTACCACG) and cpaFR (CCACCAGGTTCTTCATCGTCAGC). A fragment from gene *pleD* was amplified using primers PleDF (CCGGCATGGACGGGTTC) and PleDR (CGTTGACGCCCAGTTCCAG). A fragment from gene *dgcB* was amplified using primers DgcBF (GAGATGCTGGCGGCTGAATA) and DgcBR (CGAACTCTTCGCCACCGTAG). A fragment from gene *cleD* was amplified using primers Bresu1276F (ATCGCCGATCCGAACATGG) and Bresu1276R (TTCTCGACCCGCTTGAACAG). The fragments were then cloned into the pCR vector using the Zero Blunt cloning kit (Thermo Fisher), creating plasmids pPDC17 (*cpaF*), pLAS1 (*pleD*), pLAS2 (*dgcB*), and pLAS3 (*cleD*). These plasmids were then transformed into *B. subvibrioides* strains as previously published (3). The pCR plasmid is a non-replicating plasmid in *B. subvibrioides* that facilitates insertion of the vector into the gene of interest via recombination, thereby disrupting the gene.

To create a C-terminal *B. subvibrioides* DivJ fusion, ∼50% of the *divJ* gene covering the 3’ end was amplified by PCR using primers BSdivJgfpF (CCTCATATGGGTTTACGGGGCCTACGGG) and BsdivJgfpR (CGAGAATTCGAGACGGTCGGCGACGGTCCTG), and cloned into the pGFPC-2 plasmid (21), creating plasmid pPDC11. To create a C-terminal *B. subvibrioides* PleC fusion, ∼50% of the *pleC* gene covering the 3’ end was amplified by PCR using primers BSpleCgfpF (CAACATATGCCAGAAGGACGAGCTGAACCGC) and BspleCgfpR (TTTGAATTCGAGGCCGCCCGCGCCTGTTGTTG), and cloned into the pGFPC-2 plasmid, creating plasmid pPDC8. These plasmids are non-replicative in *B. subvibrioides* and therefore integrate into the chromosome by homologous recombination at the site of each targeted gene. The resulting integration creates a full copy of gene under the native promoter that produces a protein with C-terminal GFP tag, and a ∼50% 5’ truncated copy with no promoter. This effectively creates a strain where the tagged gene is the only functional copy.

Due to the small size of the *divK* gene, a region including the *divK* gene and ∼500 bp of sequence upstream of *divK* was amplified using primers BSdivKgfpF (AGGCATATGCCAGCGACAGGGTCTGCACC) and BsdivKgfpR (CGGGAATTCGATCCCGCCAGTACCGGAACGC) and cloned into pGFPC-2, creating plasmid pPDC27. After homologous recombination into the *B. subvibrioides* genome, two copies of the *divK* gene are produced, both under the native promoter, one of which encodes a protein C-terminally fused to GFP.

Constructs expressing *E. coli ydeH* under IPTG induction on a medium copy (pTB4) and low copy (pSA280) plasmids were originally published in (22). Constructs expressing *Pseudomonas aeruginosa pchP* under vanillate induction (pBV-5295) as well as an active site mutant (pBV-5295_E328A_) were originally published in (23).

### Transposon mutagenesis

Transposon mutagenesis was performed on the *B. subvibrioides ΔdivK* mutant using the EZ-Tn5 <KAN-2> TNP transposome (Epicentre). *B. subvibrioides ΔdivK* was grown overnight in PYE to an OD_600_ of about 0.07 [quantified with a Themo Nanodrop 2000 (Themo Scientific)]. Cells (1.5 ml) were centrifuged 15,000 x g for 3 min at room temperature. The cell pellet was then resuspended in 1 ml of water before being centrifuged again. This process was repeated. Cells were resuspended in 50 µl of nuclease free water, to which 0.2 µl of transposome was added. The mixture was incubated at room temperature for 10 minutes. The mixture was added to a Gene Pulser Cuvette with a 0.1 cm electrode gap (Bio-Rad). The cells were then electroporated as performed previously (3). Electroporation was performed using a GenePulser Xcell (Bio-Rad) at a voltage of 1,500 V, a capacitance of 25 µF, and a resistance of 400 Ω. After electroporation, cells were resuspended with 1 ml of PYE then incubated shaking at 30°C for 3 hours. Cells were diluted 3-fold then spread on PYE + Kan plates (100 µl/plate). Plates were incubated at 30°C for 5-6 days.

### Swarm assay

Strains were grown overnight in PYE, diluted to an OD_600_ of 0.02, and allowed to grow for two doublings (to OD_600_ of ∼0.06 - 0.07). All strains were diluted to OD_600_ = 0.03 and 1 µl of culture was injected into a 0.3% agar PYE plate. Isopropyl 1-thio-b-D-galactopyranoside (IPTG) (final concentration 1500 µM) and vanillate (final concentration 1 M) was added to plate mixture before pouring plates where applicable. Molten 0.3% agar in PYE (25 ml) was poured in each plate. Plates were incubated at 30°C for 5 days. Plates were imaged using a BioRad ChemiDoc MP Imaging System with Image Lab software. Swarm size was then quantified in pixels using ImageJ software. Assays were performed in triplicate and average and standard deviation were calculated.

### Short-term adhesion assay

Strains were grown overnight in PYE, diluted to an OD_600_ of 0.02, and allowed to grow for two doublings (to OD_600_ of ∼0.06 - 0.07). All strains were diluted to OD_600_ = 0.05, at which time 0.5 ml of each strain was inoculated into a well of a 24-well dish and incubated at 30°C for 2 hours in triplicate. Cell culture was removed and wells were washed 3 times with 0.5 ml of fresh PYE. To each well was added 0.5 ml of 0.1% crystal violet and incubated at room temperature for 20 minutes. Crystal violet was removed from each well before the plate was washed by dunking in a tub of deionized water. Crystal violet bound to biomass was eluted with 0.5 ml acetic acid and the A_589_ was quantified using a Themo Nanodrop 2000 (Themo Scientific). Averages for each strain were calculated and then normalized to wild-type values inoculated into the same plate. These assays were performed three times for each strain and used to calculate average and standard deviation.

### Lectin-binding assay and microscopy conditions

Holdfast staining was based on the protocol of (24). Strains of interest were grown overnight in PYE to an OD_600_ of 0.05 – 0.07. For each strain, 200 µl of culture were incubated in a centrifuge tube with 2 µl of Alexafluor 488 (GFP imaging conditions, Molecular Probes) for 20 minutes at room temperature. Cells were washed with 1 ml of sterile water then centrifuged 15,000 x g for 1 min at room temperature. The cell pellet was resuspended in 30 µl of sterile water. A 1% agarose pad (agarose in H_2_0) was prepared for each strain on a glass slide to which 1 µl of culture was added. Slides were then examined and photographed using an Olympus IX81 microscope by phase contrast and epifluorescence microscopy at appropriate wavelengths.

Holdfast of GFP-labeled strains were stained with Alexafluor 594 (RFP imaging conditions) conjugated to Wheat Germ Agglutinin and prepared for imaging as described above. Cells were imaged by phase contrast and epifluorescence microscopy at appropriate wavelengths.

### Isolation of phage

Surface water samples from freshwater bodies were collected from several sources in Lafayette County, Mississippi in 50 ml sterile centrifuge tubes and kept refrigerated. Samples were passed through 0.45 μm filters to remove debris and bacterial constituents. To isolate phage, 100 μl of filtered water was mixed with 200 μl mid-exponential *B. subvibrioides* cells and added to 2.5 ml PYE with molten 0.5% agar. The solution was poured onto PYE agar plates, allowed to harden, and then incubated at room temperature (∼22°C) for 2 days. Plaques were excised with a sterile laboratory spatula and placed into sterile 1.5 ml centrifuge tubes. 500 μl PYE was added and the sample was refrigerated overnight to extract phage particles from the agar. To build a more concentrated phage stock, the soft-agar plating was repeated with extracted particles. Instead of excising plaques, 5 ml of PYE was added to the top of the plate and refrigerated overnight. The PYE/phage solution was collected and stored in a foil-wrapped sterile glass vial, and 50 μl chloroform was added to kill residual bacterial cells. Phage solutions were stored at 4°C.

### Isolation of phage resistant mutants

*subvibrioides* was mutagenized with EZ-Tn5 transposome as described above. After electroporation, cells were grown for 3 hr without selection, followed by 3 hr with kanamycin selection. Transformed cells (100 μl) were mixed with 100 μl phage stock (∼1 x 10^10^ pfu/ml) and plated on PYE agar medium with kanamycin. Colonies arose after ∼5 days and were transferred to fresh plates. Transformants had their genomic DNA extracted using the Bactozol kit (Molecular Research Center). Identification of the transposon insertion sites was performed using Touchdown PCR (25), with transposon specific primers provided in the EZ-Tn5 kit.

### Phage sensitivity assays

Two different phage sensitivity assays were used. First (hereafter referred to as the spotting assay) involved the mixing of cells and phage in liquid suspension and then spotting droplets on an agar surface. Each cell culture was normalized to OD_600_ = 0.03. The culture was then diluted 10^-2^, 10^-4^ and 10^-5^ in PYE medium. For control assays, 5 μl of each cell suspension (including undiluted) was mixed with 5 μl PYE, then 5 μl of this mixture was spotted onto PYE plates, allowed to dry, then incubated at room temperature for 2 days. For the phage sensitivity assays, 5 μl of each cell suspension was mixed with 5 μl of phage stock (∼1 x 10^10^ pfu/ml), 5 μl spotted onto PYE plates, allowed to dry, then incubated at room temperature for 2 days.

The second assay (hereafter referred to as the soft agar assay) involved creating a lawn of cells and spotting dilutions of phage on the lawn. Cell cultures were normalized to OD_600_ = 0.03 and 200 μl of cells were mixed with 4.5 ml PYE with molten 0.5% agar, mixed, poured onto a PYE agar plate, and allowed to harden. Phage stock (∼1 x 10^10^ pfu/ml) was diluted in PYE media as individual 10X dilutions to a total of 10^-7^ dilution. 5 μl of each phage concentration (10^-1^ to 10^-7^, 7 concentrations total) were spotted on top of the soft agar surface and allowed to dry. Plates were incubated 2 days at room temperature.

### Swarm suppressor screen

Individual colonies from a transposon mutagenesis were collected on the tip of a thin sterile stick and inoculated into a 0.3% agar PYE plate. Wild-type *B. subvibrioides* strains as well as *B. subvibrioides ΔdivK* were inoculated into each plate as controls. 32 colonies were inoculated into each plate including the 2 controls. Plates were incubated at 30°C for 5 days. Plates were then examined for strains that had expanded noticeably further than the parent *divK* strain from the inoculation point. Those strains of interest were then isolated for further testing.

### Identification of swarm suppressor insertion sites

Swarm suppressor insertion sites were identified by Inverse PCR (iPCR, (26)). Genomic DNA (gDNA) was purified using the DNeasy Blood & Tissue Kit (Qiagen). Digests were then prepared using 1 µg of gDNA and either AluI or HhaI incubated overnight at 37°C. Digests were heat inactivated for 20 minutes at 80°C then column cleaned using the DNA Clean and Concentrator kit (Zymo Research). Dilute ligations (100-500 ng DNA) were then prepared so that digested fragments would likely circularize. Ligations were incubated at 17°C overnight. Reactions were heat inactivated at 65°C for 20 minutes then column cleaned using the DNA Clean and Concentrator kit. The ligated DNA was used as the template in a PCR reaction with primers that anneal inside the transposon sequence. Primers used included AluIF (GCGTTGCCAATGATGTTACAGATGAG) and AluIR (GCCCGACATTATCGCGAGCCC) as well as HhaIF2 (TTACGCTGACTTGACGGGAC) and HhaIR2 (GGAGAAAACTCACCGAGGCA). Given the large size of the resulting AluI fragment from the transposon sequence alone, another primer AluIFSeq (CGGTGAGTTTTCTCCTTCATTACAG) was designed specifically for sequencing after iPCR was complete. Primers were designed facing outward toward either end of the transposon such that the resulting PCR amplicon would be fragments that begin and end with transposon sequence with gDNA in between. PCR reactions were prepared using 10.75 µl H_2_0, 5 µl HF buffer (BioRad), 5 µl combinational enhancer solution (2.7 M betaine, 6.7 mM DTT, 6.7% DMSO, 55 µg/mL BSA), 1 µl of template DNA from each ligation, 1 µl each of their respective forward and reverse primers (primers based on what enzyme was used during digestion), 1 µl of 10 mM dNTP’s (BioLine), and 0.25 µl iProof (BioRad). PCR conditions were as follows. Initial melt was set to 98°C for 30 seconds. Melting temperature was set to 98°C for 45 seconds, annealing temperature was set to 52°C for 20 seconds, extension temperature was set to 72°C for 2:30 seconds. and these three steps were cycled through 30 times. Final extension temperature was set to 72°C for 10 minutes. 5 µl from each reaction were run on a 1% agarose gel to check for fragments. Those reactions that tested positive for bands were drop dialyzed using 0.025 µm membrane filters (Milllipore) then prepared for sequencing with their respective primers. Samples were sent to Eurofins for sequencing.

### Quantification of c-di-GMP

Strains of interest were grown overnight in PYE to an OD_600_ of 0.05 – 0.07. Metabolites were then extracted from each sample and c-di-GMP was quantified using the protocol previously described in (27). Metabolites from each strain were extracted in triplicate. Remaining cellular material was dried at room temperature and resuspended in 800 µL 0.1M NaOH. Samples were incubated at 95°C for 15 minutes. Samples were then centrifuged for 10 min at 4°C, 20,800 x g. Protein levels were measured in triplicate for each sample using 10 µl from the pellet treatment and the Pierce BCA Protein Assay Kit (Thermo Fisher Scientific). Intracellular concentrations measured by mass spectrometry were then normalized to protein levels.

## Results

### Deletion mutants in the *B. subvibrioides* DivJ-DivK-PleC system result in varied phenotypes compared to that of analogous *C. crescentus* mutations

In the previous study done in *Brevundimonas subvibrioides*, deletion mutants of the genes *divJ, divK*, and a *divJdivK* double mutant were made and partially characterized, uncovering some starkly different phenotypes compared to the homologous mutants in *C. crescentus*. However, characterization of this system was not complete as it did not extend to a key player in this system: PleC. As previously mentioned, *C. crescentus pleC* mutants display a lack of motility, pili, holdfast, and stalks (28). To begin examining the role of PleC in *B. subvibrioides*, an in-frame deletion of the *pleC* gene (Bresu_0892) was created. This strain, along with the previously published *divJ, divK*, and *divJdivK* strains, were used in a swarm assay to analyze motility. All mutant strains displayed reduced motility in swarm agar compared to the wild-type (Figure 1A). This had been reported for the published strains (3). The mechanistic reasons for this are unclear. All were observed to produce flagella and were seen to swim when observed microscopically. The *divJ* strain has significantly filamentous cell shape which is known to inhibit motility through soft agar, but the *divK* and *divJdivK* strains actually have shorter than wild-type cells. The nature of the *pleC* motility defect is also unknown. The cell size of the *pleC* mutants was not noticeably different from that of wild-type cells (Figure 1B). The *C. crescentus pleC* mutant is known to have a paralyzed flagellum which leads to a null motility phenotype, but *B. subvibrioides pleC* mutants were observed swimming under the microscope suggesting that unlike *C. crescentus* their flagellum remains functional. While the mechanistic reason for this discrepancy is unknown, it does provide another important difference in developmental signaling mutants between the two organisms.

**Figure 1.**
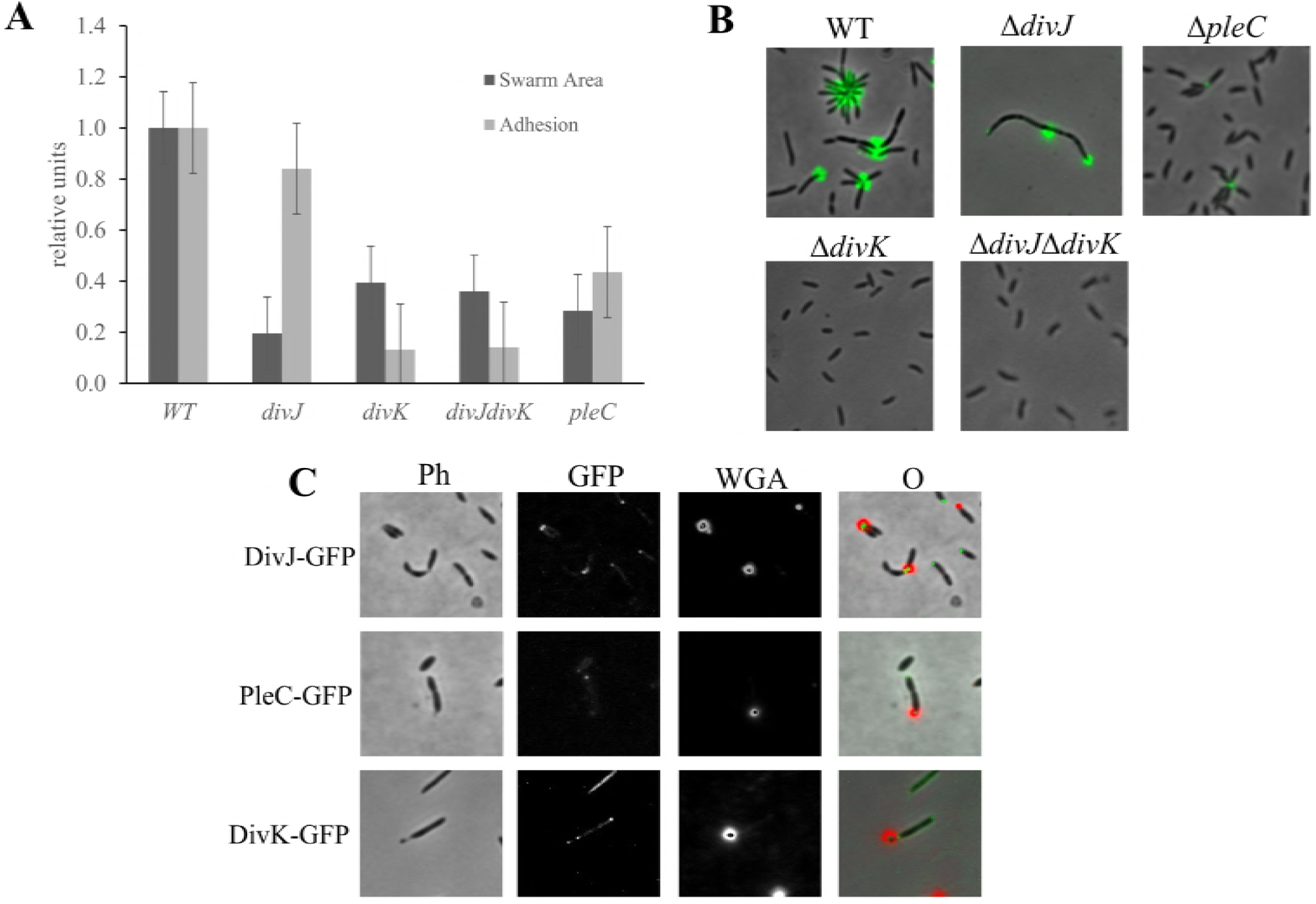
Deletions in B. subvibrioides developmental signaling genes results in varying physiological phenotypes. A) Wild-type, *divJ, divK, divJdivK*, and *pleC B. subvibrioides* strains were analyzed for swarm expansion (dark bars) and adhesion (light bars) defects using a soft agar swarm assay and a short-term adhesion assay. Mutant strains were normalized to wild-type results for both assays. Deletion of *divJ* gives motility defects but minimal adhesion defects, similar to *C. crescentus divJ* results. *B. subvibrioides divK* and *divJdivK* strains give opposite results, with severe motility and adhesion defects. The *B. subvibrioides pleC* strain has reduced motility and moderately reduced adhesion, which is similar but not identical to the *C. crescentus* Δ*pleC* strain. B) Lectin staining of holdfast material of wild-type, *divJ, divK, divJdivK*, and Δ*pleC* strains. The Δ*pleC* strain, despite having reduced adhesion in the short-term adhesion assay, still has detectable holdfast material C) GFP-tagged DivJ localizes to the holdfast producing pole, while PleC-GFP localizes to the pole opposite the holdfast. DivK-GFP displays bi-polar localization. These localization patterns are identical those of *C. crescentus* homologs.

To further the phenotypic characterization, these strains were analyzed for the surface adhesion properties using both a short-term adhesion assay as well as staining holdfast material with a fluorescently-conjugated lectin. As previously reported, the *divK* and *divJdivK* strains had minimal adhesion and no detectable holdfast material (Figure 1AB). It was previously reported that the *divJ* strain had increased adhesion over wild-type, but in this study, it was found to have slightly reduced adhesion compared to wild-type. It is not clear if this difference is significant. The *pleC* strain had reduced adhesion compared to wild-type, but more adhesion compared to the *divK* or *divJdivK* strains. When analyzed by microscopy, the *pleC* strain was found to still produce detectable holdfast, which is a difference from the *C. crescentus pleC* strain where holdfast was undetectable (28, 29).

An important component to the function of this signaling system is the subcellular localization of DivJ and PleC to the stalked and flagellar poles respectively. As the localization of these proteins had yet to be characterized in *B. subvibrioides*, GFP-tagged constructs were generated such that the tagged versions were under native expression. Because *B. subvibrioides* cells very rarely produce stalks under nutrient-replete conditions (30), holdfast material was stained using a WGA lectin conjugated with a fluorophore that uses RFP imaging conditions. As seen in Figure 1C, DivJ-GFP formed foci at the same pole as the holdfast, while PleC-GFP formed foci at poles opposite holdfast. As it has been demonstrated that holdfast material is produced at the same pole as the stalk in *B. subvibrioides* (30), this result suggests that these proteins demonstrate the same localization patterns as their *C. crescentus* counterparts. Additionally, DivK-GFP was seen to form bipolar foci in predivisional cells (Figure 1C), the same as *C. crescentus* DivK. Therefore, while the phenotypic consequences of signaling protein disruption vary between these organisms, the localization patterns of the proteins are consistent.

### Isolation of a bacteriophage capable of infecting *B. subvibrioides*

Another important developmental event in *C. crescentus* is the production of pili at the flagellar pole coincident with cell division. Pili are very difficult to visualize, and in *C. crescentus* the production of pili in strains of interest can be assessed with the use of the bacteriophage ΦCbK, which infects the cell using the pilus. Resistance to the phage indicates the absence of pili. However, bacteriophage that infect *C. crescentus* do not infect *B. subvibrioides* (data not shown) despite their close relation. In an attempt to develop a similar tool for *B. subvibrioides*, a phage capable of infecting this organism was isolated.

Despite the fact that *B. subvibrioides* was isolated from a freshwater pond in California over 50 years ago (4), a phage capable of infecting the bacterium was isolated from a freshwater pond in Lafayette County, Mississippi. This result is a testament to the ubiquitous nature of *Caulobacter* and *Brevundimonas* species in freshwater environments all over the globe. This phage has been named Delta, after the state’s famous Mississippi Delta region. To determine the host range for this phage, it was tested against multiple *Brevundimonas* species (Figure 2A). Delta has a relatively narrow host range, causing the largest reduction of cell viability in *B. subvibrioides* and *B. aveniformis*, with some reduction in *B. basaltis* and *B. halotolerans* as well. None of the other 14 *Brevundimonas* species showed any significant reduction in cell viability. Neither did Delta show any infectivity toward *C. crescentus* (data not shown). While *B. subvibrioides, B. aveniformis*, and *B. basaltis* all belong to the same sub-clade within the *Brevundimonas* genus (P. Caccamo, Y.V. Brun, personal communication), so do *B. kwangchunensis, B. alba* and *B. lenta*, all of which are more closely related to *B. subvibrioides* than *B. aveniformis* and all of which were resistant to the phage. Therefore, infectivity does not appear to fall along clear phylogenetic lines and may be determined by some other factor.

**Figure 2.**
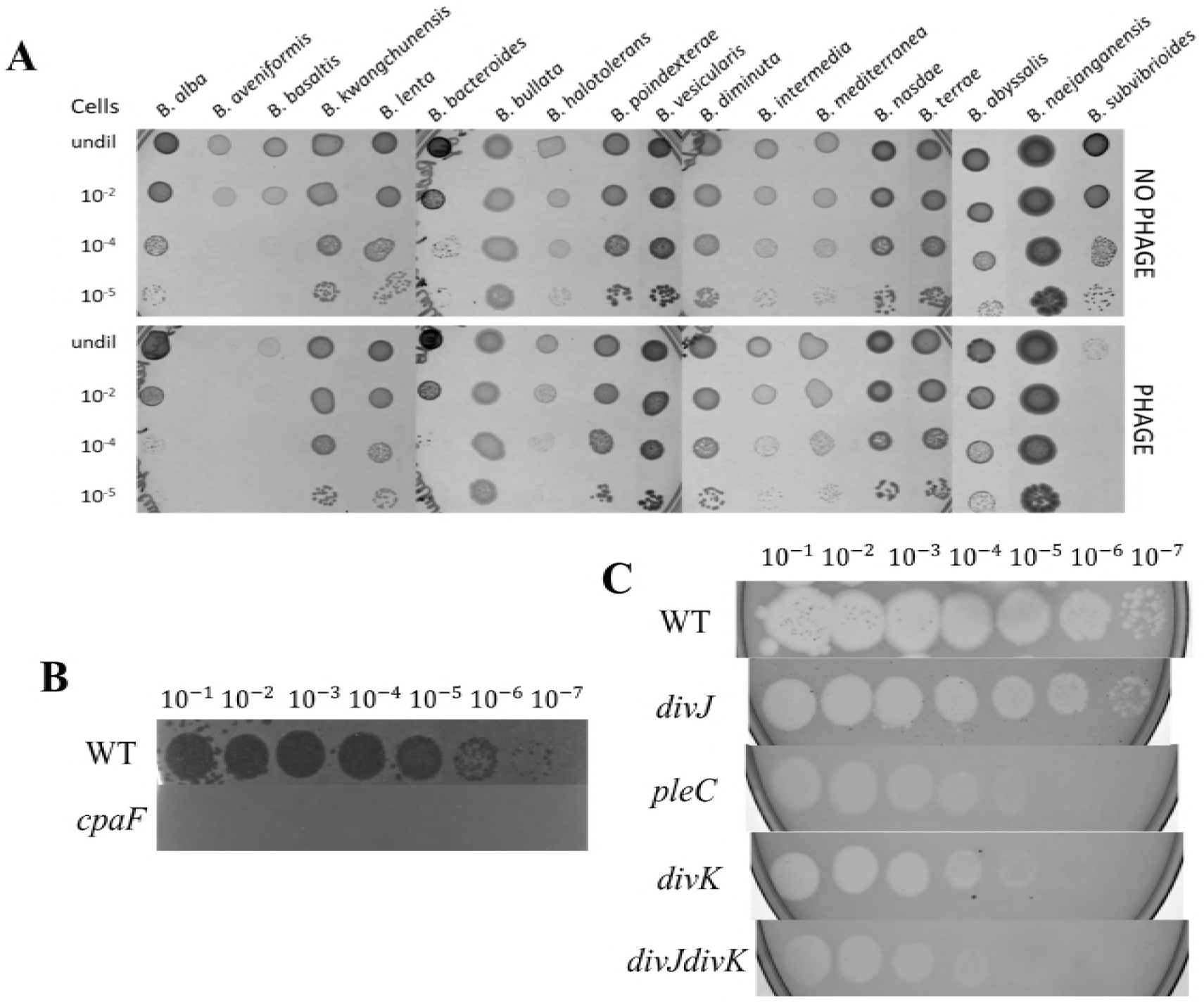
Bacteriophage Delta serves as a tool to investigate *B. subvibrioides* pilus production. A) Phage Delta was tested for infection in 18 different *Brevundimonas* species. Control assays used PYE media instead of phage stock. Delta caused a significant reduction in *subvibrioides* and *B. aveniformis* viability, with some reduction in *B. basaltis* and *B. halotolerans* as well. B) Phage Delta was tested against wild-type and *cpaF*∷pCR *B. subvibrioides* strains using a soft agar phage assay. Wild-type displayed zones of clearing with phage dilutions up to 10^-7^, while the *cpaF* strain showed resistance to all phage dilutions. C) *B. subvibrioides* developmental signaling mutants were tested with phage Delta in soft agar phage assays. Wild-type shows clear susceptibility to Delta, as does the *divJ* strain suggesting that, like *crescentus divJ*, it produces pili. The *pleC* strain shows a 2-3 orders of magnitude reduced susceptibility to the phage, indicating reduced pilus production which is consistent with the *C. crescentus* phenotype. The *divK* and *divJdivK* strains display similar to resistance as the *pleC* strain. Here again, *divK* disruption causes the opposite phenotype to *divJ* disruption, unlike the *C. crescentus* results.

To begin identifying the infection mechanism of Delta, *B. subvibrioides* was randomly mutagenized with a Tn5 transposon and resulting transformants were mixed with Delta to select for transposon insertions conferring phage resistance as a way to identify the phage infection mechanism. Phage resistant mutants were readily obtained and maintained phage resistance when rescreened. A number of transposon insertion sites were sequenced and several were found in the pilus biogenesis cluster homologous to the *C. crescentus flp*-type pilus cluster. Insertions were found in the homologs for *cpaD, cpaE* and *cpaF*; it is known disruption of *cpaE* in *C*. *crescentus* abolishes pilus formation and leads to ΦCbK resistance (2, 3, 31–33). A targeted disruption was made in *cpaF* and tested for phage sensitivity by the soft agar assay (Figure 2B). The *cpaF* disruption caused complete resistance to the phage. The fact that multiple transposon insertions were found in the pilus cluster and that the *cpaF* disruption leads to phage resistance strongly suggest that Delta utilizes the *B. subvibrioides* pilus as part of its infection mechanism. The identification of another pili-tropic phage is not surprising as pili are major phage targets in multiple organisms.

Phage Delta was used to assess the potential pilis production in developmental signaling mutants using the soft agar assay (Figure 2C). The *divJ* mutant has similar susceptibility to Delta as the wild-type, suggesting this strain still produces pili. This result is consistent with the *C. crescentus* result as the *C. crescentus divJ* mutant is ΦCbK susceptible (8). Conversely, the *B. subvibrioides pleC* mutant shows a clear reduction in susceptibility to Delta, indicating that this strain is deficient in pilus production. If so, this would also be consistent with the *C. crescentus pleC* mutant which is resistant to ΦCbK (8, 28). With regards to the *divK* strain, if that mutant were to follow the *C. crescentus* model it should demonstrate the same susceptibility as the *divJ* strain. Alternatively, as the *divK* strain has often demonstrated opposite phenotypes to *divJ* in *B. subvibrioides*, one might predict it to demonstrate resistance to Delta. As seen in Figure 2C, the *divK* strain (and the *divJdivK* strain) shows the same level of resistance to phage Delta as the *pleC* mutant. Therefore, in regards to phage sensitivity, the *divK* strain is once again opposite of the prediction of the *C. crescentus* model. Interestingly, none of these developmental signaling mutants demonstrate complete resistance to Delta as seen in the *cpaF* strain. This result suggests that these mutations impact pilus synthesis, but does not abolish it completely.

### A suppressor screen identifies mutations related to c-di-GMP signaling

As the *B. subvibrioides divK* mutant displays the most unusual phenotypes with regard to the *C. crescentus* model, this strain was selected for further analysis. Complementation of *divK* was attempted by expressing wild-type DivK from an inducible promoter on a replicating plasmid, however induction failed to complement any of the *divK* phenotypes (data not shown), indicating proper complementation conditions have not yet been identified. Transposon mutagenesis was performed on this strain and mutants were screened for those that restore motility. Two mutants were found (Bresu_1276 and Bresu_2169) that restored motility to the *divK* strain, and maintained this phenotype when recreated by plasmid insertional disruption. Both mutants were involved in c-di-GMP signaling. The *C. crescentus* homolog of the Bresu_1276 gene, CC3100 (42% identical to Bresu_1276), was recently characterized in a subcluster of CheY-like response regulators and renamed CleD (34). Function of CleD is, at least in part, initiated by binding c-di-GMP via an arginine-rich residue with high affinity and specificity for c-di-GMP (34). Upon binding, roughly 30% of CleD localizes to the flagellated pole of the swarmer cell. Nesper et. al suggests that CleD may bind directly to the flagellar motor switch protein, FliM. Based upon these findings, it was hypothesized that increased c-di-GMP levels cause activation of CleD, which binds to the flagellar switch and inhibits flagellar function. In *C. crescentus, cleD* mutants are 150% more motile while their adhesion does not differ significantly from that of the wild-type. Unlike conventional response regulators, the phosphoryl-receiving aspartate is replaced with a glutamate in CleD. In other response regulators, replacement of the aspartate with a glutamate mimics the phosphorylated state and locks the protein in an active conformation. Alignment of CleD with orthologs from various *Caulobacter* and *Brevundimonas* species demonstrated that this was a conserved feature of CleD within this clade (Figure 3). Similar to *C. crescentus*, the swarm size of *B. subvibrioides cleD* mutant increased to 151% compared to wild-type. However, unlike *C. crescentus* the *cleD* disruption reduced adhesion by 35% compared to wild-type (Figure 4A). A knockout of *cleD* in the *divK* background led to a complete restoration of motility compared to that of wild-type, while adhesion did not appear to be affected. These phenotypes correspond relatively well with the model given in Nesper et al. A cell lacking CleD would have decreased flagellar motor disruption leading to an increase in motility and a delay in surface attachment.

**Figure 3.**
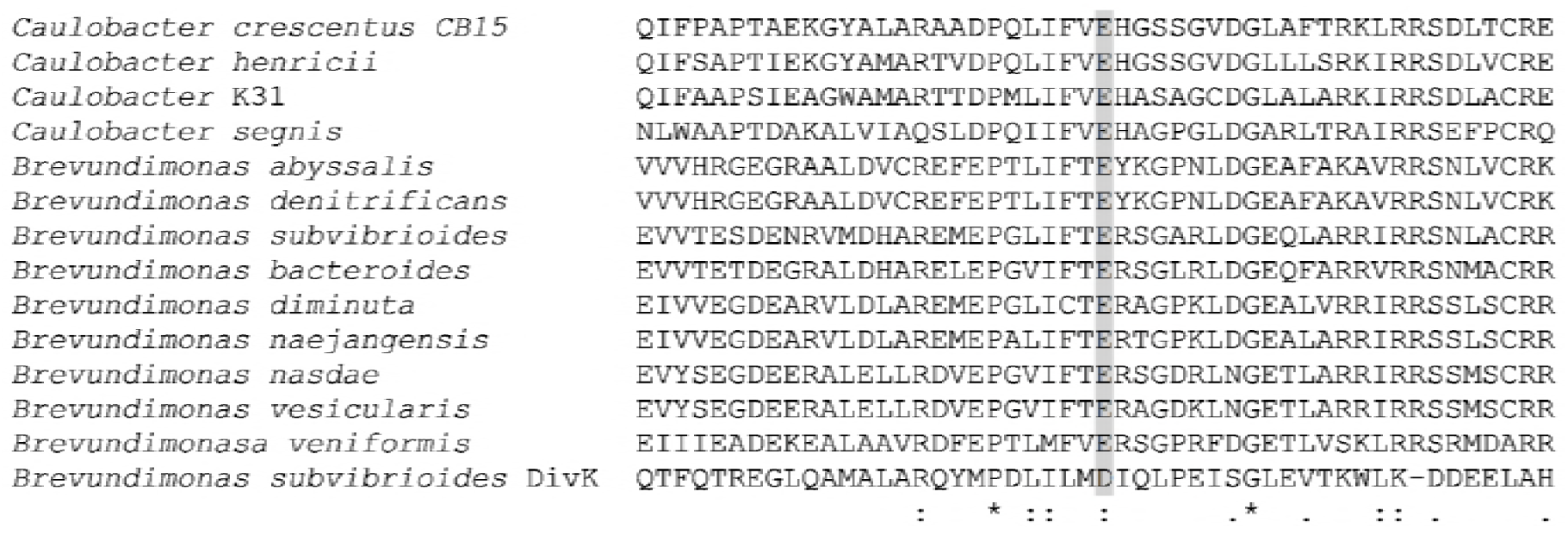
CleD displays a conserved glutamate residue in place of an aspartate typical of response regulators. CleD orthologs from various Caulobacter and Brevundimonas species were aligned by ClustalW, along with B. subvibrioides DivK. The shaded box indicates *B. subvibrioides* DivK D53, which is analogous to *C. crescentus* DivK D53 and is the known phosphoryl-accepting residue. This alignment demonstrates that CleD orthologs all contain a glutamate substitution at that site, which has been found to mimic the phosphorylated state and lock the protein in an active conformation in other response regulators.

**Figure 4.**
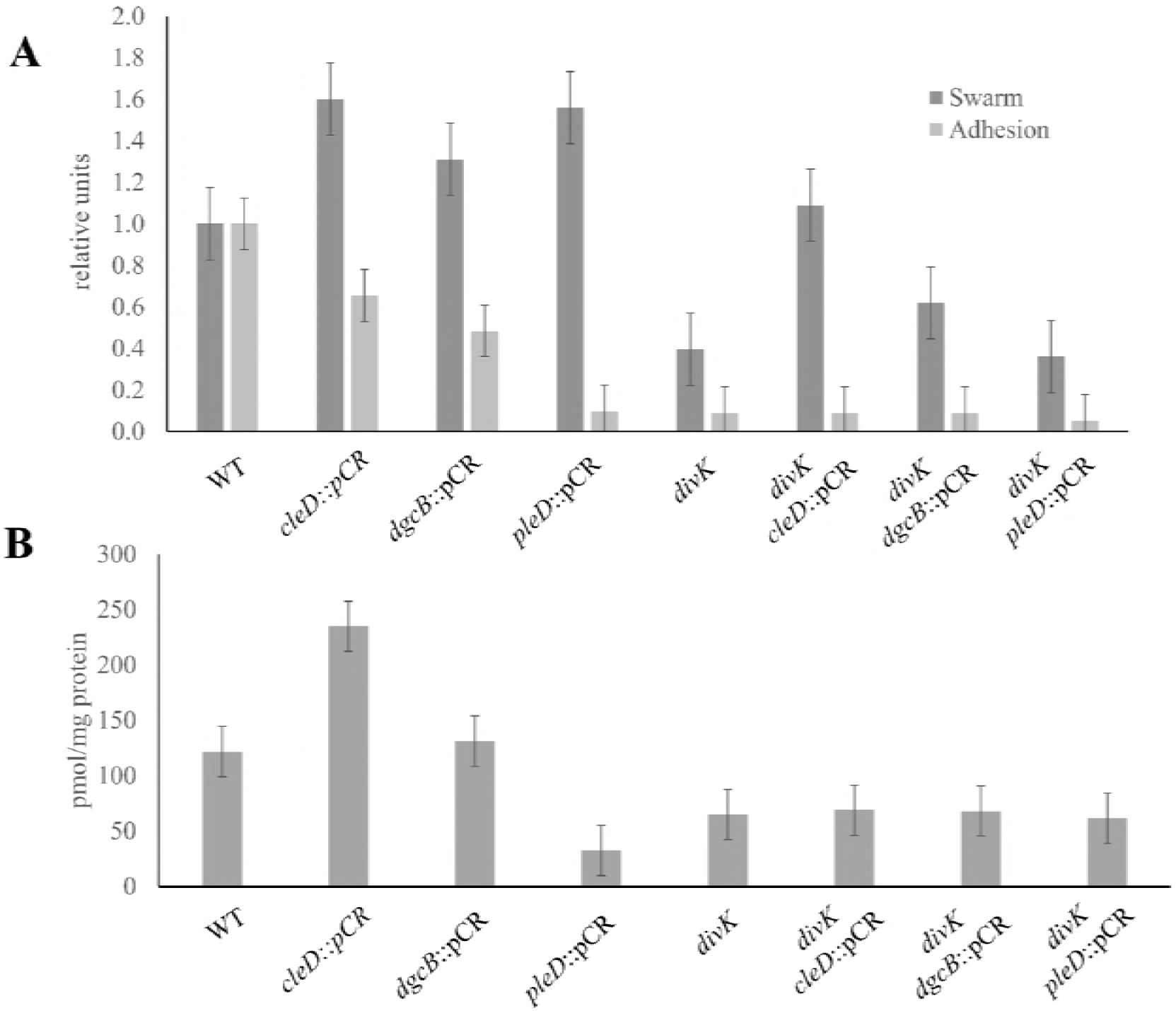
Phenotypes exhibited by *divK* suppressors do not coincide with intracellular c-di-GMP levels. A) Swarm expansion (dark bars) and surface adhesion (light bars) of suppressor mutations tested in both the wild-type and *divK* background. Disruption of CleD, DgcB and PleD lead to increased motility in the wild-type background, but only CleD and DgcB lead to increased motility in the *divK* background. Disruptions in the wild-type background lead to varying levels of adhesion reduction, but the same disruptions had no effect on adhesion in the *divK* background. B) C-di-GMP levels were measured using mass spectrometry then normalized to the amount of biomass from each sample. Despite disruptions causing increased motility in the wild-type background, those strains had different c-di-GMP levels. No disruption changed c-di-GMP levels in the *divK* background even though some strains suppressed the motility defect while others did not. These results show a discrepancy between phenotypic effects and intracellular c-di-GMP levels.

Bresu_2169 is the homolog of the well-characterized *C. crescentus* diguanylate cyclase, DgcB (61% identical amino acid sequence). In *C. crescentus*, DgcB is one of two major diguanylate cyclases that work in conjunction to elevate c-di-GMP levels which in turn helps regulate the cell cycle, specifically in regards to polar morphogenesis (35). It has been shown that a *dgcB* mutant causes adhesion to drop to nearly 50% compared to wild-type while motility was elevated to almost 150%. It was unsurprising to find very similar changes in phenotypes in the *dgcB* mutant in wild-type *B. subvibrioides*. In the *dgcB* mutant, swarm expansion increased by 124% while adhesion dropped to only 46% compared to wild-type (Figure 4A). Though the *dgcB* mutant did not restore motility to wild-type levels in the *divK* background, the insertion did cause the swarm to expand nearly twice as much as that of the *divK* parent. These phenotypes are consistent with our current understanding of c-di-GMP’s role in the *C. crescentus* cell cycle. As c-di-GMP builds up in the cell, it begins to make the switch from its motile phase to its sessile phase. Deleting a diguanylate cyclase therefore should prolong the swarmer cell stage, thereby increasing motility and decreasing adhesion.

### A *pleD* mutant lacks hypermotility in *divK* background

Given the identification of *dgcB* in the suppressor screen, it was of note that the screen did not identify the other well-characterized diguanylate cycle involved in the *C. crescentus* cell cycle, PleD. PleD is an atypical response regulator with two receiver domains in addition to the diguanylate cyclase domain (36, 37). The *pleD* mutant in *C. crescentus* has been shown to suppresses the *pleC* motility defect in *C. crescentus* which led to its initial discovery alongside *divK* (22, 36, 37). However, in a wild-type background, *pleD* disruption has actually been shown to reduce motility to about 60% compared to wild-type (22, 35). Additionally, a 70% reduction in adhesion is observed in *pleD* mutants which is thought to be a result of delayed holdfast production (22, 35, 38). Therefore, it was not clear whether a *pleD* disruption would lead to motility defect suppression in a *divK* background. To examine this, a *pleD* disruption was made in both the wild-type and *divK B. subvibrioides* strains (Figure 4A).

In wild-type *B. subvibrioides, pleD* disruption caused hypermotility with swarms expanding to 156% of wild-type, while adhesion dropped to only 10% compared to wild-type. While this data supports the broader theory of c-di-GMP’s role as the “switch” between the motile and sessile phase of the cell cycle, it does not align with those phenotypes seen in a *C. crescentus pleD* mutant. While adhesion is reduced in both organisms, the reduction in adhesion was much more drastic in *B. subvibrioides* than *C. crescentus*. Moreover, the motility phenotypes in homologous *pleD* mutants shift in opposite directions. In *C. crescentus, pleD* mutants causes a decrease in motility by nearly 40% in the wild-type background (22, 35). In *B*. *subvibrioides*, we see a 156% increase (Figure 4A).

Another interesting detail discovered in performing these assays was the lack of change in phenotypes seen in the *pleD* disruption strain in a *divK* background. It is not surprising that adhesion was not negatively impacted as it is already significantly lower in the *divK* strain compared to wild-type. However, disrupting the *pleD* gene did not cause hypermotility in the *divK* mutant even though it does cause hypermotility in the wild-type background. In fact, motility was reduced to 89% compared to the *divK* control (Figure 4A). It is not clear why disruption of the diguanylate cyclase DgcB leads to increased motility in both the wild-type and *divK* backgrounds, but disruption of another diguanylate cycle PleD leads to increased motility in just the wild-type background. Interestingly, it was previously shown that DivJ and PleC do not act on DivK alone, but in fact also have the same enzymatic functions on PleD phosphorylation as well (39). It may be that PleD acts upon motility not through c-di-GMP signaling but instead by modulating DivK activity, perhaps by interacting/interfering with the polar kinases. If so, then the absence of DivK could block this effect.

### Suppressor mutants have altered c-di-GMP levels

As these mutations are all involved in c-di-GMP signaling, c-di-GMP levels in each strain were quantified to determine if the cellular levels in each strain correspond to observed phenotypes. These metabolites were quantified from whole cell lysates. In bacteria, high c-di-GMP levels typically induce adhesion while low c-di-GMP levels induce motility. Therefore, it would be expected that hypermotile strains would show decreased c-di-GMP levels. Instead, hypermotile strains of the wild-type background had varying c-di-GMP levels (Figure 4B). The *pleD* knockout had reduced c-di-GMP levels as predicted. While it may seem surprising that c-di-GMP levels are not affected in a *dgcB* mutant, this in fact true of the *C. crescentus* mutant as well (35). This result suggests that the c-di-GMP levels found in the *dgcB* strain do not appear to be the cause for the observed changes in motility and adhesion.

Perhaps the most interesting result is that the *cleD* mutant had the highest c-di-GMP levels of all strains tested. This is surprising as it is suggested by Nesper et. al. that CleD does not affect c-di-GMP levels at all, but rather is affected by them. CleD is a response regulator that contains neither a GGDEF nor an EAL domain characteristic of diguanylate cyclases and phosphodiesterases respectively. Instead it is thought CleD binds to c-di-GMP, which then stimulates it to interact with the flagellar motor. The data presented here suggests that there may be a feedback loop whereby increased motility in the swarm agar leads to increased c-di-GMP levels. One potential explanation is that this situation increases contact with surfaces. Yet the *cleD* mutant clearly shows decreased adhesion compared to wild-type despite the elevated c-di-GMP levels. Therefore, there must be a block between the high c-di-GMP levels and the execution of those levels into adhesion in this strain.

Very different results were obtained when c-di-GMP levels were measured in *divK* derived strains (Figure 4B). While a wide variety of motility phenotypes were observed in *cleD, dgcB*, and *pleD* disruptions in the *divK* background, their c-di-GMP levels are all nearly identical to that of the *divK* mutant. For the *dgcB divK* strain, once again the increase in motility occurs without a change in c-di-GMP levels. These results suggest that DgcB is not a significant contributor to c-di-GMP production in *B. subvibrioides*. While *pleD* disruption leads to decreased c-di-GMP levels in the wild-type background, no change is seen in the *divK* background. This means in the absence of PleD some other enzyme must be responsible for achieving these levels of c-di-GMP. Given the lack of impact DgcB seems to have on c-di-GMP signaling, it is tempting to speculate an as-yet characterized diguanylate cyclase is involved. Lastly the elevated c-di-GMP levels seen in the *cleD* disruption are not seen when *cleD* is disrupted in the *divK* background. This result suggests that whatever feedback mechanism leads to elevated c-di-GMP levels is not functional in the *divK* mutant.

### Non-native diguanylate cyclases and phosphodiesterases cause shifts in c-di-GMP levels but do not alter phenotypes in the *divK* strain

As previously mentioned, c-di-GMP is thought to assist in the coordination of certain developmental processes throughout the cell cycle. The previous results found mutations in genes involved in c-di-GMP signaling could suppress developmental defects, but the actual effect of the mutations appears uncoupled from effects on c-di-GMP levels. In order to further investigate the connection between developmental defects and c-di-GMP signaling, c-di-GMP levels were artificially manipulated. Plasmid constructs expressing non-native c-di-GMP metabolizing enzymes previously used in similar experiments in *C. crescentus* were obtained and expressed in *B. subvibrioides*. The diguanylate cyclase *ydeH* from *Escherichia coli* was expressed from two different IPTG inducible plasmids; a medium copy number pBBR-based plamid pTB4, and a low copy number pRK2-based plasmid, pSA280 (22). The combination of the two different inducible copy number plasmids resulted in different elevated levels of c-di-GMP (Figure 5B). A phosphodiesterase *pchP* from *Pseudomonas aeruginosa* (40), as well as its active site mutant *pchP*^E^328^A^ were expressed from pBV-MCS4, a vanillate inducible medium copy number plasmid (23). The phosphodiesterase on a medium copy plasmid was enough to decrease levels of c-di-GMP to either equivalent or lower levels as is seen in the *divK* strain. The decrease was not observed when the active site mutant was expressed, demonstrating that the reduction of c-di-GMP was the result of *pchP* expression. Wild-type and *divK* strains were grown with IPTG and vanillate respectively to control for any growth effects caused by the inducers.

**Figure 5.**
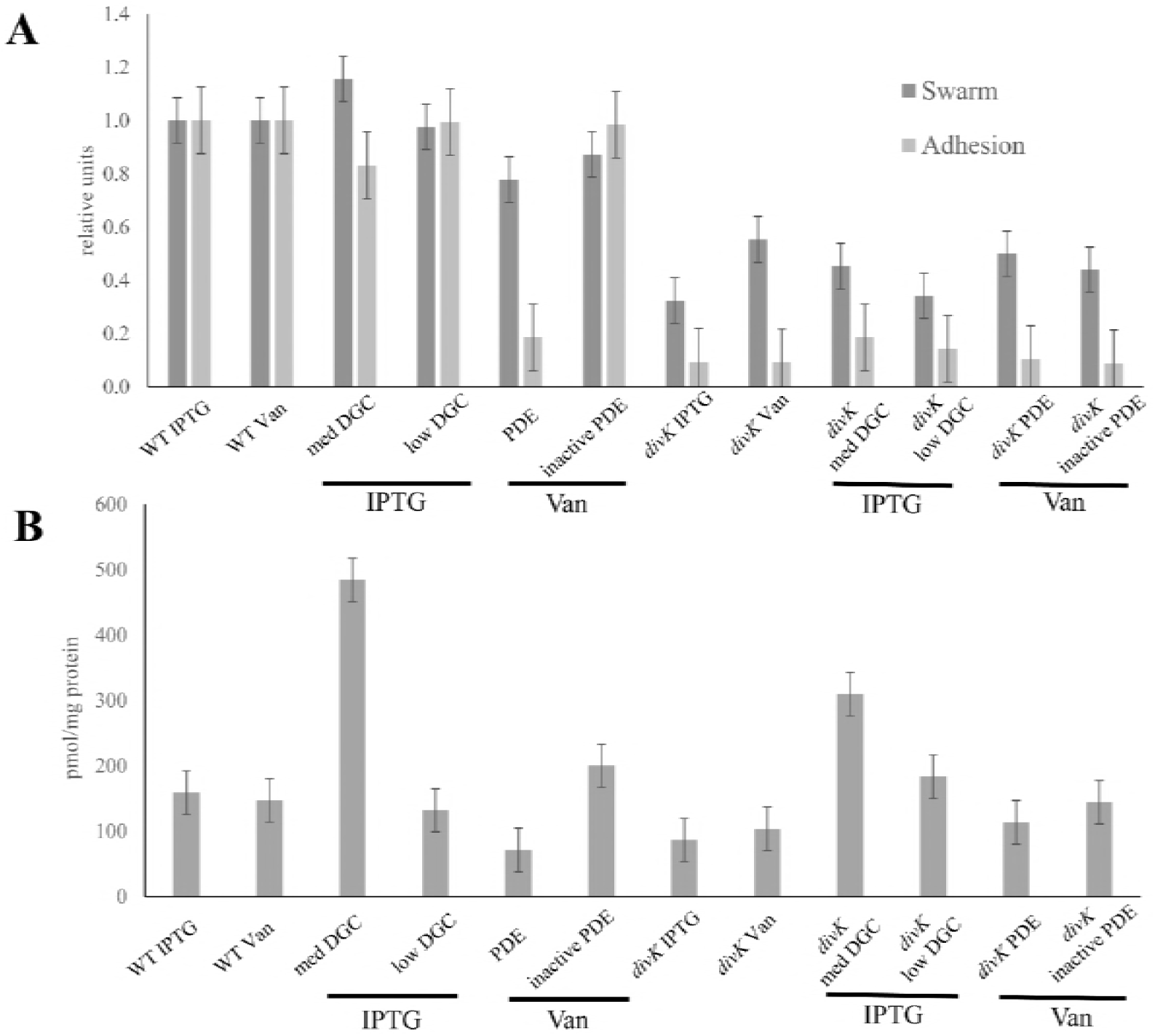
Artificial manipulation of c-di-GMP levels do not significantly affect phenotypes in the *divK* mutant. A) Swarm expansion (dark bars) and surface adhesion (light bars) of strains that have altered c-di-GMP levels caused by expression of non-native enzymes in the wild-type and *divK* background. Constructs including the *E. coli* diguanylate cyclase *ydeH* expressed from a medium copy plasmid (med DGC) and a low copy plasmid (low DGC), the *P. aeruginosa* phosphodiesterase *pchP* (PDE) as well as a catalytically inactive variant (inactive PDE). Bars below the x-axis outline inducer used for plasmids in each strain. In the wild-type background the medium copy DGC increased motility and decreased adhesion, which is opposite the expected outcome, while the PDE reduced motility and severely reduced adhesion. In the *divK* background, no expression construct significantly altered the phenotypes. B) C-di-GMP levels were measured using mass spectrometry then normalized to the amount of biomass from each sample. In the wild-type background the medium copy DGC significantly increased c-di-GMP levels while the PDE reduced c-di-GMP levels. In the *divK* background, both DGC constructs increased c-di-GMP levels, though PDE expression has no effect, despite the fact that neither DGC construct has an effect on motility and adhesion phenotypes.

The low copy diguanylate cyclase plasmid did not appear to affect levels (Figure 5B), and unsurprisingly did not appear to affect either motility or adhesion. However, the medium copy diguanylate cyclase plasmid increased c-di-GMP levels but had the opposite phenotypic effect than expected. An increase in c-di-GMP levels would be predicted to decrease motility and increase adhesion, but here the increase in c-di-GMP caused an increase in motility and a decrease in adhesion (Figure 5A). Conversely, expression of the phosphodiesterase in the wild-type background caused a reduction in c-di-GMP which would be predicted to increase motility and decrease adhesion. While this strain had a large reduction of adhesion, it also had a reduction in motility. Therefore, the changes in c-di-GMP levels largely do not match the changes in phenotype. It is also interesting to note that expression of the phosphodiesterase results in similar c-di-GMP levels to that of the *divK* strain yet the phosphodiesterase strain demonstrates much larger swarm sizes than the *divK* strain.

In the *divK* background strain, expression of either diguanylate cyclase increases c-di-GMP levels, though the low copy diguanylate cyclase increase is not as dramatic as the medium copy. However, neither expression level has a significant impact on motility or adhesion (Figure 5). Neither the phosphodiesterase nor its active site mutant cause a noticeable shift in the c-di-GMP levels compared to the *divK* strain nor any noticeable impact on phenotype. In fact, though the c-di-GMP levels differed dramatically between strains, the phenotypes of all six of these strains are not impacted. T-tests performed between each strain and its respective control showed no significant difference. These results appear to be the antithesis of those found from the suppressor screen. While the suppressor mutants showed recovery in their motility defect compared to *divK*, their c-di-GMP levels did not significantly differ from each other or *divK*. Conversely, when c-di-GMP levels were artificially manipulated, alterations of c-di-GMP levels in the *divK* strain had no impact on phenotypes. These results suggest that DivK is somehow serving as a block or a buffer to c-di-GMP levels and their effects on phenotypes and calls into question the role c-di-GMP has in *B. subvibrioides* developmental progression.

## Discussion

Across closely related bacterial species, high levels of gene conservation are commonly observed. It has therefore been a long-standing assumption that information gathered from studying a model organism can be extrapolated to other closely related organisms. Through this study, it has been shown that these assumptions may not be as safe to make as previously thought. Preliminary data raised a few questions by demonstrating major differences in the phenotypes of *divK* mutants between species. Here more differences between systems were observed by demonstrating *pleC* mutants have similar but not identical phenotypes to *C. crescentus pleC* mutants. These differences in signaling protein operation were observed despite the fact that subcellular localization patterns are the same in both organisms. These findings strongly indicated that some aspect of this system is somehow behaving differently than the understood *C. crescentus* model system. These discoveries not only raised questions about how the DivJ-DivK-PleC system has evolved over a short evolutionary distance, but they have also called into question different aspects of the *C. crescentus* system.

In an attempt to further map this system in *B. subvibrioides* and perhaps identify missing pieces, a suppressor screen was employed using the *divK* mutant as its phenotypes differed most dramatically from its *C. crescentus* homolog. Suppressor mutations were found in genes predicted to encode proteins that affected or were affected by c-di-GMP. This was not necessarily a surprising discovery. C-di-GMP is a second messenger signaling system conserved across many bacterial species used to coordinate the switch between motile and sessile lifestyles. Previous research in *C. crescentus* suggests that organism integrated c-di-GMP signaling into the swarmer-to-stalked cell transition. Mutations that modify c-di-GMP signaling would be predicted to impact the swarmer cell stage, perhaps lengthening the amount of time the cell stays in that stage and thus lead to an increase in swarm spreading in soft agar. However, further inquiry into c-di-GMP levels of *divK* suppressor mutants revealed discrepancies between c-di-GMP levels and their corresponding phenotypes. Firstly, CleD, a CheY-like response regulator that is thought to affect flagellar motor function, caused the strongest suppression of the *divK* mutant restoring motility levels to that of wild-type. Given that the reported function of CleD is to bind the FliM filament of the flagellar motor and interfere with motor function to boost rapid surface attachment (34), it is expected that disruption of *cleD* would result in increased motility and decreased adhesion which can be seen in both the wild-type and *divK* background strains (Figure 4A). What was unexpected, however, was to find that a lack of CleD led to one of the highest detected levels of c-di-GMP in this study, which was surprising given that CleD has no predicted diguanylate cyclase or phosphodiesterase domains. Yet when this same mutation was placed in the *divK* background, the c-di-GMP levels were indistinguishable from the *divK* parent. Therefore, the same mutation leads to hypermotility in two different backgrounds despite the fact that c-di-GMP levels are drastically different. Consequently, the phenotypic results of the mutation do not match the c-di-GMP levels, suggesting that c-di-GMP has little or no effect on the motility phenotype. A similar result was seen with DgcB. Disruption of *dgcB* in either the wild-type or *divK* background resulted in hypermotility, but c-di-GMP levels were not altered. Once again, the effect on motility occurred independently of c-di-GMP levels.

Disruption of *pleD* led to different results. In the wild-type background, disruption of *pleD* caused a reduction in c-di-GMP levels, which would be predicted given that PleD contains a diguanylate cyclase domain. Therefore, of the three proteins analyzed here (CleD, DgcB, and PleD), it appears only PleD actually contributes to the c-di-GMP pool. Yet when disruptions of any of the genes are placed in a *divK* background, c-di-GMP levels are not altered. Disruption of *divK* seems to somehow stabilize c-di-GMP levels. Even when non-native enzymes are expressed in the *divK* background the magnitude of changes seen in the c-di-GMP pool is dampened compared to the magnitude of change seen when the enzymes are expressed in the wild-type background. This may explain why *pleD* was not found in the suppressor screen. While CleD and DgcB seem involved in c-di-GMP signaling, their effect on the cell appears c-di-GMP-independent, while PleD appears to perform its action by affecting the c-di-GMP pool. If that pool is stabilized in the *divK* strain, then disruption of *pleD* will have no effect on either the c-di-GMP pool or on the motility phenotype. However, it should be noted that c-di-GMP levels were measured from whole cell lysates and does not reflect the possibility of spatial variations of c-di-GMP levels within the cell. It is possible that CleD and DgcB have c-di-GMP dependent effects, but those effects are limited to specific sub-cellular locations in the cell.

This research raises several questions. First, what is the exact role of c-di-GMP in cell cycle progression of *B. subvibrioides*? Is this signal a major driver of the swarmer cell and swarmer cell differentiation? Or have the various c-di-GMP signaling components found new roles in the swarmer cell and the actual c-di-GMP is simply vestigial. What is the role of PleD in cell cycle progression? Why are c-di-GMP levels so stable when DivK is removed? And lastly, are the answers to these questions specific to *B. subvibrioides*, or can they be extrapolated back to *C. crescentus*? Further investigation into c-di-GMP signaling in both organisms is required.

## Acknowledgements

The authors wish to thank P. Caccamo and Y.V. Brun for providing most of the *Brevundimonas* species, as well as the iPCR protocol that had been adapted for use in *C. crescentus*, U. Jenal for providing several plasmids, Satish Adhikari and Kari Ferolito for their help with cloning, and the Curtis lab for general support. This work was supported by the United States National Science Foundation CAREER program award 1552647.

